# APOE4 increases energy metabolism in APOE-isogenic iPSC-derived neurons

**DOI:** 10.1101/2024.06.03.597106

**Authors:** Vanessa Budny, Yannic Knöpfli, Debora Meier, Kathrin Zürcher, Chantal Bodenmann, Siri L. Peter, Terry Müller, Marie Tardy, Cedric Cortijo, Christian Tackenberg

## Abstract

The apolipoprotein E4 (APOE4) allele represents the major genetic risk factor for Alzheimer’s disease (AD). In contrast, APOE2 is known to lower the AD risk while APOE3 is defined as risk neutral. APOE plays a prominent role in the bioenergetic homeostasis of the brain, and early-stage metabolic changes have been detected in brains of AD patients. Although APOE is primarily expressed by astrocytes in the brain, neurons also have been shown as source for APOE. However, little is known about the differential role of the three APOE isoforms for neuronal energy homeostasis. In this study, we generated pure human neurons (iN cells) from APOE-isogenic induced pluripotent stem cells (iPSCs), expressing either APOE2, APOE3, APOE4 or carrying an APOE-knockout (KO) to investigate APOE isoform-specific effects on neuronal energy metabolism. We showed that endogenously produced APOE4 enhanced mitochondrial ATP production in APOE-isogenic iN cells but not in the corresponding iPS cell line. This effect neither correlated with the expression levels of mitochondrial fission or fusion proteins, nor with the intracellular or secreted levels of APOE, which were similar for APOE2, APOE3 and APOE4 iN cells. ATP production and basal respiration in APOE-KO iN cells strongly differed from APOE4 and more closely resembled APOE2 and APOE3 iN cells indicating a gain-of-function mechanism of APOE4 rather than a loss-of-function. Taken together, our findings in APOE isogenic iN cells reveal an APOE genotype-dependent and neuron-specific regulation of oxidative energy metabolism.

## Introduction

Alzheimer’s Disease (AD) is the most common age-related neurodegenerative disorder affecting about 50 million people world-wide [1]. While disease-causing mutations are very rare, genome-wide association studies have identified 90 independent genetic variants associated with AD susceptibility [2]. Among them, the APOE4 allele represents the strongest risk factor. APOE has three major isoforms, APOE2, APOE3 and APOE4, which differ only in two amino acid residues. While APOE3/3 is the most common genotype and is defined as risk-neutral, the APOE2 allele is protective but occurs only in 10 % of AD patients and 20 % of healthy controls. In contrast, the presence of one or two copies of APOE4 increases the risk by three- or twelve-fold, respectively [3]. In the brain, APOE is mainly expressed by astrocytes, but also microglia and neurons have been shown to secrete APOE. Studies on the selective removal of APOE from the mouse brain indicated that astrocytic APOE makes up 75-80 % of total brain APOE while neurons contribute 15-20% [4]. Astrocytes secrete approx. 50-60 % of APOE they produce, while neurons retain most APOE intracellularly and only secrete approx. 10 % [5,6] indicating significant differences in APOE biology between neurons and astrocytes. However, if and how neuronal APOE mediated AD pathophysiology remains unclear [7].

APOE has a prominent role in the bioenergetic homeostasis of the brain. The high energy demand of the brain renders it sensitive to changes in energy supply and alterations in the consumption of glucose as well as deficits in mitochondrial functions are hallmarks of AD [8]. Postmortem studies suggest that reduced brain glucose utilization occurs in patients suffering from AD [9]. Further, FDG-PET analyzes in AD patients revealed that cortical glucose hypometabolism is particularly pronounced in APOE4 carriers [10].

Several studies have been conducted on how APOE4 affects energy metabolism in cell lines or murine primary cultures with partially contradictory results. However, it is also important to use human cell models to better understand these mechanisms as cellular functions or expression of several disease-relevant proteins, including APOE, may strongly differ between rodent and human neuronal cells [11]. The use of isogenic lines thereby allows the distinct analysis of APOE effects without bias caused by interpatient genetic variation. Further, we still lack the mechanistic insights into how different APOE isoforms – not only the risk-increasing APOE4 but also the protective APOE2 – affect the energy metabolism in human neurons.

In the present study, we show that endogenously produced APOE4 increases mitochondrial energy metabolism in pure APOE-isogenic human iPSC-derived neurons (iN cells) but not in the respective iPS cell line. This effect was independent of APOE levels in the different cell lines and did not correlate to levels of mitochondrial fission or fusion proteins. ATP production, as well as basal respiration of APOE-KO iN cells were comparable to APOE2 and strongly differed from APOE4 iN cells indicating a gain-of-function mechanism of APOE4 rather than a loss-of-function.

## Materials and Methods

### iPS cell culture

APOE-isogenic iPS cell lines BIONi010-C3 (APOE-KO), BIONi010-C6 (APOE2), BIONi010-C2 (APOE3), and BIONi010-C4 (APOE4) [12–14] have been purchased from the European Bank of induced pluripotent Stem Cells (EBiSC). iPSCs were cultured on vitronectin (1:25) (100-0763, StemCell Technologies) coated plates in mTESR+ medium (100-0276, StemCell Technologies), split every 3-4 days and culture medium was exchanged every other day. For splitting, cells were washed with DPBS, incubated with ReLeaSR (5872, StemCell Technologies) for 4 min at 37 °C with 5 % CO_2_, detached in 1 mL mTESR+ and transferred to a new plate.

### iN cell differentiation

For iPSC differentiation into iN cells we used the overexpression of Ngn2 as previously described, with slight modification [15,16]. On day minus one, 500’000 cells per well were seeded as single cells onto new 6-well plates (6 WPs) coated with vitronectin and cultured in mTESR+ medium supplemented with thiazovivin (SML1045, Sigma Aldrich). 2 μL of each viral construct (TetO-Ngn2-P2A-puromycin, rtTA, TetO-EGFP) was added to every well. Lentiviral production was performed as previously described [15]. On day zero, the medium was replaced by neural induction medium (N2 1:100, non-essential amino acids (NEAA) 1:100 (11140050, Thermo Fisher), doxycycline 2 mg/L (D9891, Sigma), BDNF 10 ng/mL (450-02, Peprotech), NT3 10 ng/mL (450-03, Peprotech), laminin 0.2 μg/mL (L2020, Sigma) in DMEM/F12). 24 h later, the medium was replaced by fresh medium supplemented with 0.5 mg/L puromycin (P9620, Sigma Aldrich) to select for transduced cells. On day two, cells were replated to 6 WPs to produce protein or RNA samples, onto 65 % nitric acid pretreated coverslips in a 24 WP for immunocytochemistry (ICC), or to Seahorse XF24 V7 PS plates (100777-004, Agilent) for seahorse assays. Plates were coated with 100 μg/mL Poly-L-Lysin (P8920, Sigma Aldrich) and 3.4 μg/mL Laminin (L2020, Sigma Aldrich). Cells were detached with Accutase (A6964, Sigma Aldrich) and seeded as single cells (750’000 for 6WP, 100’000 for 24WP) onto the PLL/Laminin coated plates with neuronal differentiation medium (B27 supplement 1:50 (17504001, Gibco), Glutamax 1 mM, doxycycline 2 mg/L, BDNF 10 ng/mL, NT3 10 ng/mL, GDNF 10 ng/mL (450-10, Peprotech), CNTF 10 ng/mL (450-13, Peprotech), laminin 0.2 μg/mL, cAMP 0.5 mM (Cay18820-500, Biomol) in neurobasal medium) and thiazovivin. On day three, the full medium was exchanged to remove thiazovivin. On day five, more medium was gently added (1 mL for 6 WP, 500 μL for 24 WP). Starting from day six, only half of the medium was removed and replaced by fresh medium (1500 μL for 6 WP, 500 μL for 24 WP). 2 μM AraC (C1768, Sigma) was added to the medium from day six until the end of the differentiation. Half the medium was exchanged every 3-4 days until day 21. All samples were taken, and experiments were performed on day 21 or 22.

### Immunocytochemistry

iPSCs were fixed one day after replating. iN cells were fixed after differentiation at d21 for 20 minutes at room temperature (RT) with 4 % Paraformaldehyde (PFA) (47377.9L, VWR) and 4 % sucrose (S9378, Merck) in DPBS. After washing the cells three times with PBS for about 5 minutes at RT, cells were blocked with 10 % donkey serum (D9663, Sigma Aldrich) and 0.1 % Triton (X100, Sigma Aldrich) in DPBS for 1 h at RT. After washing with DPBS, cells were incubated with primary antibodies diluted in 3 % donkey serum and 0.1 % Triton in DPBS overnight at 4 °C. After washing at the next day, secondary antibodies diluted in 3 % donkey serum and 0.1 % Triton in DPBS were added to the cells and incubated for 2 h at RT in the dark. After washing, cells were incubated with 0.4 ng/μl DAPI (D9542, Sigma Aldrich) diluted in DPBS for 10 minutes. After another washing step, the coverslips were mounted with Mowiol (81381, Sigma Aldrich) on microscope objectives and stored over night at 4 °C protected from light.

| *Primary antibody* | *Producer* | *Cat. No.* | *Dilution* |
| --- | --- | --- | --- |
| Anti-Oct4 | Cell Signaling | 2890S | 1:200 |
| Anti-Nanog | Thermo Fisher | 14-5768-82 | 1:100 |
| Anti-MAP2 | Synaptic systems | 188011 | 1:500 |

| *Secondary antibody* | *Producer* | *Cat. No.* | *Dilution* |
| --- | --- | --- | --- |
| Dk-α-ms-cy3 | Jackson ImmunoResearch | 715-165-151 | 1:1000 |
| Dk-α-rb-cy5 | Jackson ImmunoResearch | 715-175-152 | 1:1000 |
| Dk-α-rb-Alexa647 | Jackson ImmunoResearch | 715-605-152 | 1:250 |

### RNA extraction

Cells were harvested with Accutase. 1 Mio cells were resuspended in 350 μL RLT buffer and 3.5 μL β-mercaptoethanol (31350-010, Gibco). RNA extraction was performed by using the RNeasy Mini Kit (74104, Qiagen) according to the manufacturer’s instructions. RNA concentrations of the samples were analyzed with the NanoDrop spectrophotometer (Thermo Fisher Scientific) and stored at -80 °C.

### Protein extraction

iPSCs were harvested with Accutase, while iN cells were harvested using as cell scraper and resuspended in RIPA buffer supplemented with protease inhibitor (11697498001, Sigma Aldrich). Due to high viscosity of iPSC samples, 2 μL Benzonase (E1014, Merck) in 1 mL sample was added. Four cycles of 30s of sonication were used to disrupt the cellular membranes. To extract the proteins, samples were centrifuged at 20’000 g for 10 min at 4 °C and the supernatant was finally collected and stored at -20 °C until usage. Protein concentrations (μg/μL) were determined with the Pierce BCA Assay Kit (23252, Thermo Fisher Scientific) according to the manufacturer’s instructions and the absorption at 562 nm was measured with the Infinite M Nano plate reader (Tecan).

### Meso Scale Discovery (MSD) immunoassay for APOE

The APOE MSD assay (K151AMLR-2, MSD) was performed according to the standard assay protocol 1 from the manufacturer. The plate was coated with 25 μL of biotinylated capture antibody pre-diluted in coating diluent, incubated for 1 h shaking at RT and washed three times with washing buffer. A serial dilution of the standard was prepared by diluting the 20x stock calibrator in assay diluent, resulting in a standard curve ranging from 750’000 pg/mL to 0 pg/mL. Samples were diluted as following: iN cell lysates undiluted, iN cell supernatant undiluted, iPSC lysates 1:50 and iPSC supernatant 1:3. 1.5 mL supernatant samples of iN cells was lyophilized before and resuspended in 100 μL RIPA. 30 μL of the calibrator standard or sample was added to the coated plate (in duplicate). The plate was incubated shaking for 1 h at RT. After washing 3x, 50 μL of detection antibody solution was added to each well and again incubated shaking for 1 h at RT. After washing 3x, 150 μL of read buffer was added to each well and the plate was immediately analyzed on an MSD instrument. By using the absorbance of the standard, a standard curve was determined and used to calculate the total APOE levels of the tested samples in relation to the respective dilutions. Measured APOE levels were normalized to total protein concentrations (μg/μL) (*see protein extraction*).

### Western blotting

Each sample was diluted in RIPA buffer to obtain a concentration of 10 μg and mixed with sample buffer (NP0007, Thermo Fisher Scientific). Samples were denatured for 5 min at 95 °C. Seeblue2 plus protein ladder (LC5925, Thermo Fisher Scientific) and samples were loaded onto 10-20 % Tricine SDS-PAGE gels (EC6625BOX, Invitrogen) and run at 60 V for 15 min and 100 V for 90 min. Blotting was performed with the Trans-Blot Turbo Mini 0.2 μm nitrocellulose Transfer Pack (1704158, Bio-Rad) and the Trans-Blot Turbo Transfer System (1704158, Bio-Rad) at 2.5 A with 25 V for 7 min. Membranes were washed with 0.05 % Tween (P1379, Sigma Aldrich) in PBS and blocked with 5 % milk solution (A0830, ITW Reagents) in PBS for 1 h at RT shaking. Membranes were then washed three times with PBS-Tween and incubated with primary antibodies diluted in 5 % milk solution in PBS-Tween overnight at 4 °C shaking. The next day, membranes were washed three times with PBS-Tween and incubated with secondary antibodies diluted in 5 % milk solution in PBS-Tween for 2 h at RT in the dark shaking. Membranes were washed three times, developed with one of the ECL selection kits (RPN2232/ RPN2235, Cytiva; 32106, Thermo Fisher Scientific) and imaged at the Image Quant 800 (Cytiva). Background subtraction was performed, and protein expressions were normalized to GAPDH.

Antibodies used for western blot.
| *Primary antibody* | *Producer* | *Cat. No.* | *Dilution* |
| --- | --- | --- | --- |
| Anti-MFN1 | Cell Signaling | 14739S | 1:250 |
| Anti-MFN2 | Cell Signaling | 11925S | 1:1000 |
| Anti-FIS1 | Abcam | ab15686 | 1:10'000 |
| Anti-OPA1 | Cell Signaling | 67589S | 1:2000 |
| Anti-GAPDH | Meridian | H86504M | 1:5000 |

| *Secondary antibody* | *Producer* | *Cat. No.* | *Dilution* |
| --- | --- | --- | --- |
| Dk-α-ms-peroxidase | Jackson ImmunoResearch | 715-035-151 | 1:5000 |
| Dk-α-rb-peroxidase | Jackson ImmunoResearch | 111-035-144 | 1:5000 |

### qRT-PCR

Cell-type specific markers MAP2, TUBB3, OCT4 and mitochondrial fusion/ fission markers MFN1, MFN2, OPA1 and FIS1 were analyzed. For details on the primers see Table S1. Data was normalized to the housekeeping marker GAPDH. The detection method was SYBR green (1725121, Bio-Rad). For each sample 5 μL SYBR green, 0.05 μL forward primer, 0.05 μL reverse primer, 2 μL sample, and 2.9 μL nuclease-free water were used. The qRT-PCR protocol was as follows: Separation of the cDNA in an initial hold stage of 10 min at 95°C, followed by 40 amplification cycles of 15s at 95 °C and 1 min at 60 °C and finally one melting curve cycle of 15s at 95 °C, 1 min 60 °C and 15s at 95 °C.

### Seahorse assay

Using the Seahorse XFe24 (Agilent) the oxygen consumption rate (OCR), the extracellular acidification rate (ECAR) and the proton efflux rate (PER) were measured. Based on these measurements, all other factors were calculated. Seahorse experiments were performed according to the manufacturer’s instructions. Sensors were preincubated in H_2_O at 37 °C without CO_2_ overnight and changed to calibrant 1 h before the assay. Optimal cell density for each cell type was tested in prior Seahorse experiments. iPSCs were plated at 40’000 cells/well the day before the assay. iN cells were replated at 100’000 cells/well at day 2 of the iN differentiation protocol and analyzed at day 21 or day 22. For each assay four measurements at baseline and three measurements after drug induction were performed. To avoid detachment of iN cells during the Mito Stress Test, each condition was shorted by one measurement, resulting in three baseline measurements and two measurements after each drug. Each assay type was repeated three times for each cell line and included four background wells for each assay. The Seahorse DMEM medium (103575-100, Agilent) was freshly supplemented with 10 mM glucose, 1 mM pyruvate and 2 mM L-glutamine. For the ATP Rate Assay drugs were used at the final concentrations of 1.5 μM oligomycin and 0.5 μM Rotenone + Antimycin A (ROT/AA). For the Mito Stress Test drugs were used at the final concentrations of 1.5 μM oligomycin, 1 μM Carbonyl cyanide-4 phenylhydrazone (FCCP) and 0.5 μM Rotenone + Antimycin A (ROT/AA). FCCP titration was done in earlier Seahorse experiments. After Seahorse assays, cells were immediately fixated with 4 % PFA and 4 % sucrose in DPBS and stained with 0.4 ng/μL DAPI for 10 min at RT. DAPI was imaged with an inverted fluorescence microscope (Zeiss) with 10x magnification. The number of nuclei in each well was automatically analyzed by a macro written in Fiji and used for normalization of the Seahorse data with the Wave Software.

### Data analysis

Normal distribution was tested by using the Shapiro-Wilk normality test and the Kolmogorov-Smirnov test. Outliers were identified with the ROUT test (Q = 1 %). Differences between more than two groups were either analyzed by one-way ANOVA for normally distributed data or by the Kruskal-Wallis test for not normally distributed data. One-way ANOVA was followed by a Tukey test for multiple comparisons. A Kruskal-Wallis test was always followed by a Dunn’s multiple comparisons test. A p value of less than 0.05 was considered significant.

## Results

### APOE-isogenic iPSCs differentiate into iN cells and express similar amounts of APOE

APOE-isogenic iPS cell lines BIONi010-C3 (APOE-KO), BIONi010-C6 (APOE2), BIONi010-C2 (APOE3), and BIONi010-C4 (APOE4) were used for differentiation into iN cells (Fig. 1A). iPSC pluripotency was confirmed by immunostaining for OCT4 and NANOG (Fig. 1B). iN cell differentiation was achieved by lentiviral overexpression of Neurogenin 2 (Ngn2) [15,16]. iN cells were co-transfected with EGFP to monitor their differentiation status based on cell morphology (Fig. S1), showing a network of differentiated neurons at day 21. Mature iN cells were positive for MAP2 demonstrating the successful neuronal differentiation (Fig. 1C). No differences were observed in marker expression and differentiation potential between the four APOE-isogenic lines (Fig. 1B, C). qPCR confirmed that iPSCs were positive for OCT4 while iN cells show upregulation of MAP2 and βIII-tubulin (TUBB3) mRNA (Fig. 1D). APOE was detected in APOE2, -E3 and -E4 iPSCs and iN cells (Fig. 1E, F) while no APOE protein was present in APOE-KO cells. APOE levels did not significantly differ between the APOE lines. In both cell types, iPSCs and iN cells, APOE was mainly local-ized intracellularly (85-92 % intracellular in iPSCs; 96-98 % intracellular in iN cells) with overall higher APOE levels per total protein in iPSCs compared to iN cells.

**Figure 1:**
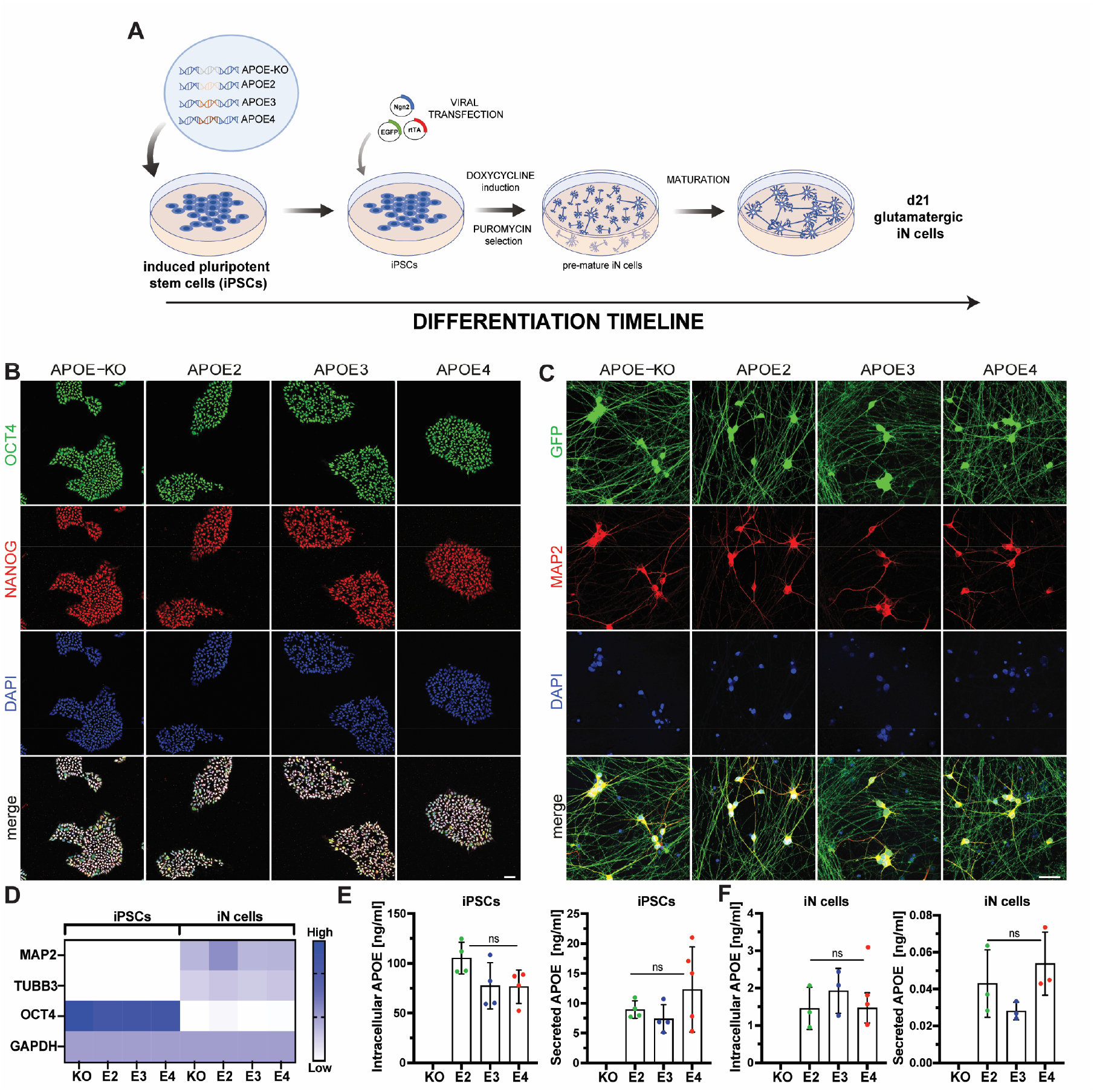
Differentiation of APOE-isogenic iPSCs into iN cells. (**A**) Schematic timeline of APOE-isogenic iN cell differentiation. (**B**) Representative confocal images of OCT4, NANOG and DAPI in isogenic iPSCs. (**C**) Representative images of GFP, MAP2 and DAPI in isogenic iN cells. (**D**) Relative gene expression of cell specific iPSC cell (OCT4) and neuronal marker (MAP2 and βIII-tubulin (TUBB3) in iPSCs and iN cells, measured by qPCR. (**E, F**) Intracellular and secreted APOE protein levels normalized to total protein concentration in iPSCs and in iN cells, measured by MSD. E-F: n=4 independent samples, one-way ANOVA with Tukey multiple comparison test. Scale bars: 50 μm.

### APOE4 iN cells show higher mito and glyco ATP production than APOE3, -E2 and -KO cells

APOE has been associated with altered brain energy metabolism, while the mechanism and the differential effect of the three major APOE isoforms are still largely unknown. To determine how the APOE genotype affects glycolytic and mitochondrial respiration-based ATP production in human cells, a Seahorse ATP rate assay was performed. Oxygen consumption rate (OCR) and extracellular acidification rate (ECAR) were measured four times at baseline and three times after each drug administration in all APOE lines in iPSCs (Fig. 2A, B) and iN cells (Fig. 2C, D). Oligomycin and Rotenone/Antimycin A were used to inhibit respiratory complex V and I/III, respectively, based on which mitochondrial ATP production could be analyzed. Both, APOE4 iPSCs and iN cells displayed the highest glycoATP production while no difference was observed between APOE2, -E3 and -KO (Fig. 2E, I). No APOE genotype effect was found on mitochondrial ATP production in iPSCs (Fig. 2F). In contrast, APOE4 iN cells had significantly higher mitochondrial ATP production compared to iN cells from the other APOE lines (Fig. 2J) suggesting a cell type-specific APOE4 effect. Total ATP production, i.e. the sum of mitoATP and glycolATP, was highest in APOE4 cells in both cell types, iPSCs and iN cells (Fig. 2G, K). However, total ATP production in APOE4 iPSCs did only significantly differ from APOE2 iPSCs, while APOE4 iN cells showed significantly higher total ATP production compared to all other iN cell lines. (Fig. 2G, K). As expected, iPSCs gained most of their energy through glycolysis (approx. 60 % of total ATP is glycoATP) (Fig. 1H), whereas iN cells mainly relied on oxidative phosphorylation (approx. 70 % of total ATP is mitoATP) (Fig. 1L).

**Figure 2:**
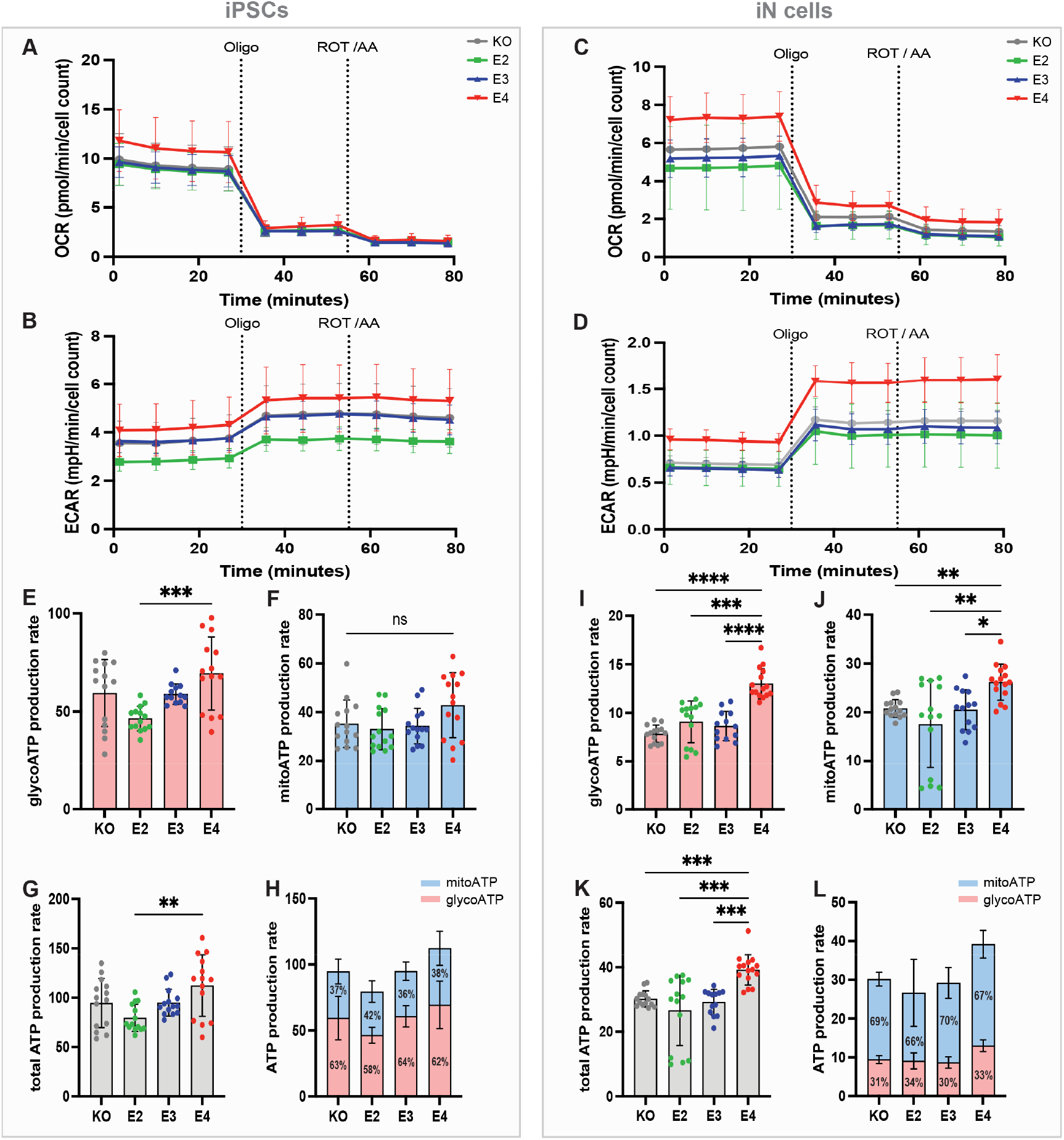
Seahorse ATP rate assay in APOE-isogenic iPSCs and iN cells. (**A**) Oxygen consumption rate (OCR) and (**B**) extracellular acidification rate (ECAR) in iPSCs. (**C**) OCR and (**D**) ECAR in iN cells. APOE-KO in grey, APOE2 in green, APOE3 in blue and APOE4 in red. Oligo: Oligomycin; ROT/AA: Rotenone/Antimycin A. (**E**) Glycolytic ATP (glycoATP) production rate, (**F**) Mitochondrial ATP (mitoATP) production rate and (**G-H**) total ATP production rate in iPSCs. (**I**) GlycoATP production rate, (**J**) mitoATP production rate (**K-L**) total ATP production rate in iN cells. GlycoATP is shown in red and mitoATP in blue. Kruskal-Wallis test (E-H, J-L) and one-way ANOVA (I). ns = not significant, * p < 0.05, ** p < 0.01, *** p < 0.001, **** p < 0.0001.

### APOE genotype does not affect levels of mitochondrial fission and fusion proteins in iN cells

A key for efficient energy production is maintaining a functional and healthy mitochon-drial network. This is ensured by dynamic reshaping events called fusion and fission (Fig. 3A). To investigate whether APOE genotype influences these processes in iPSCs and/or iN cells, mitochondrial proteins involved in fusion and fission were analyzed (Fig. 3B). No differences between the APOE lines were observed for the mitochondrial fusion proteins mitofusin-1/2 (MFN1/2) and OPA1, in iPSCs (Fig. 3C, D, F) or iN cells (Fig. 3G-J). Mito-chondrial fission marker FIS1 was decreased in APOE4 iPSCs compared to E2 (Fig. 3E), but not changed in iN cells (Fig. 3I).

**Figure 3:**
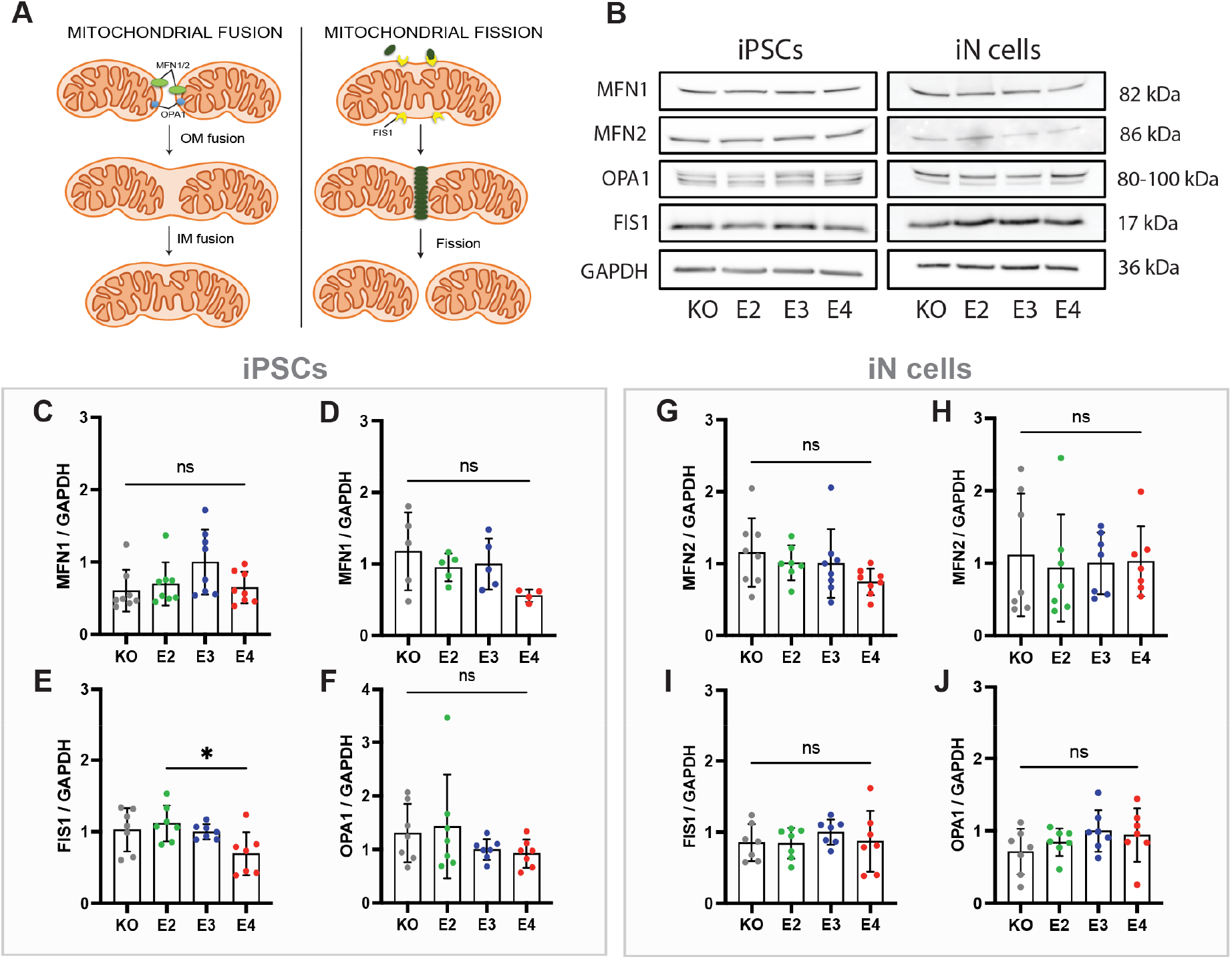
Levels of mitochondrial fusion and fission proteins. (**A**) Schematic overview of mitochondrial fusion and fission markers. (**B**) Representative western blot images of MFN1, MFN2, FIS1, OPA1 and GAPDH in APOE-KO, -E2, -E3 and -E4 isogenic iPSCs and iN cells. (**C-F**) Quantified protein levels of MFN1, MFN2, FIS1 and OPA1 in APOE-KO, -E2, -E3 and-E4 isogenic iPSCs. (**G-J**) Quantified protein levels of MFN1, MFN2, FIS1 and OPA1 in APOE-KO, -E2, -E3 and -E4 isogenic iN cells. All protein levels have been normalized to GAPDH. Kruskal-Wallis test (C, D, F) and one-way ANOVA (E, G-J). ns = not significant, * p < 0.05, ** p < 0.01, *** p < 0.001. OM: outer mitochondrial membrane; IM: inner mitochondrial membrane

### APOE regulates mitochondrial respiration and respiratory capacity in a genotype-dependent manner

As we observed an APOE genotype effect on mitochondrial ATP production specifically in iN cells but not in iPSCs (Fig. 2F, J), we investigated mitochondrial function in more detail. Basal and maximal respiration as well as mitochondrial capacity and proton leak were analyzed using Seahorse Mitochondrial Stress Test. OCR and ECAR were measured at baseline and after the addition of Oligomycin, carbonyl cyanide-4 (trifluoromethoxy) phenylhydrazone (FCCP) and ROT/AA in iPSCs (Fig. 4A, B) and iN cells (Fig. 4C, D). OCR values were used to calculate mitochondrial properties. To induce stress, FCCP was administered which leads to the collapse of the proton gradient and disruption of the mitochondrial membrane potential resulting in an uninhibited electron flow through the electron transport chain and maximum oxygen consumption when reaching complex IV. In agreement with our results from the ATP rate assay (Fig. 2F), iPSCs lines did not differ in basal and maximal mitochondrial respiration (Fig. 4E, F). In contrast, APOE4 iN cells showed the highest basal as well maximal respiration (Fig. 4I, J) which aligns with the higher mitoATP production shown above (Fig. 2J). It should be noted that significant differences were only observed between APOE4 and APOE-KO iN cells. Spare respiratory capacity is an indicator of how well cells can respond to an energetic demand reflecting cell fitness or flexibility. This was found to be highest in APOE2 iPSCs and iN cells (Fig. 4G, K) compared to the other APOE genotypes of the respective cell type. No significant changes in proton leak were observed in any cell line (Fig. 4H, L). Taken together, APOE4 increased mitochondrial respiration in iN cells but not in iPSCs.

**Figure 4:**
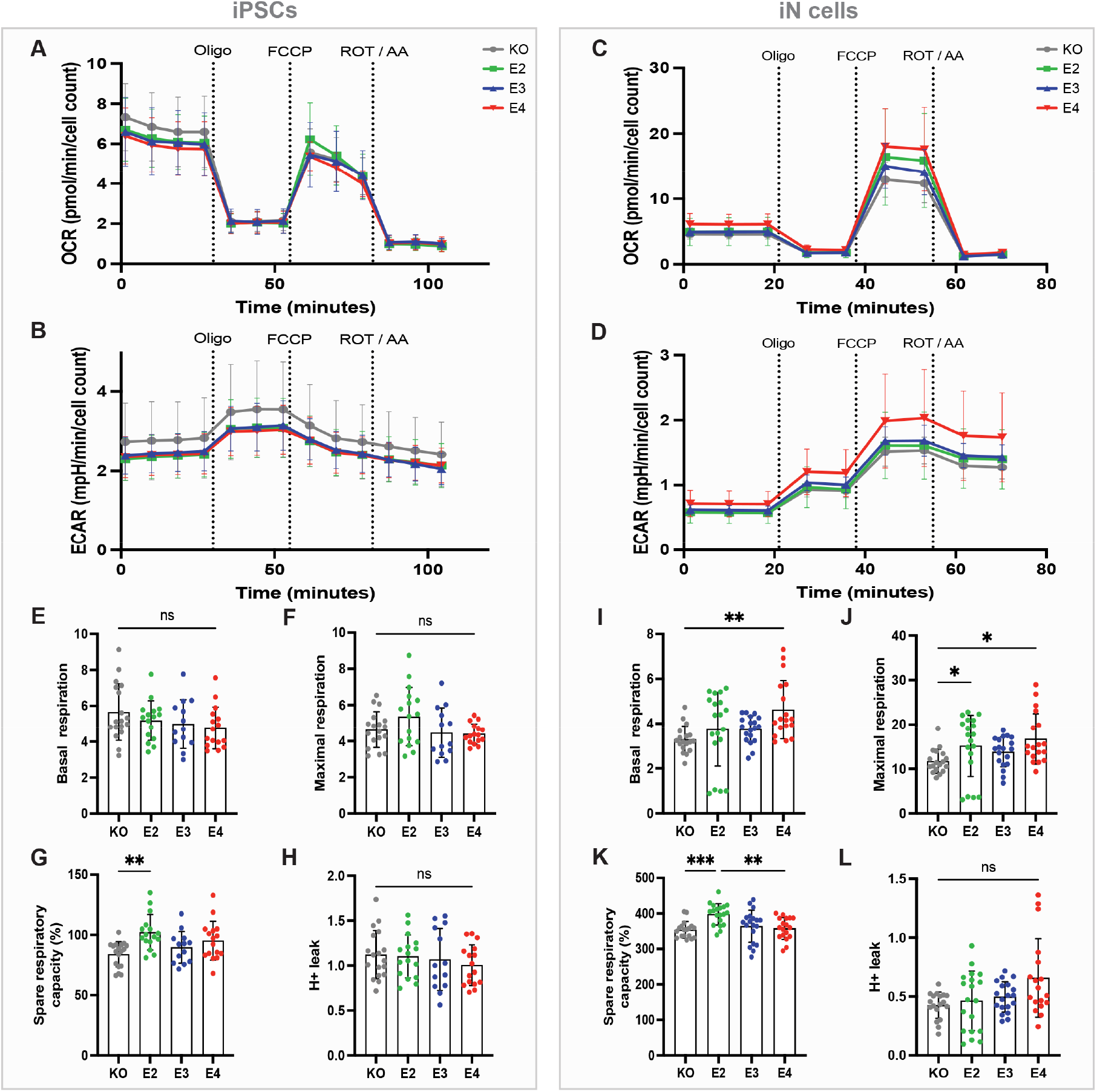
Seahorse Mitochondrial Stress Test in APOE-isogenic iPSCs and iN cells. (**A**) Oxygen consumption rate (OCR) and (**B**) extracellular acidification rate (ECAR) in iPSCs. (**C**) OCR and (**D**) ECAR in iN cells. APOE-KO in grey, APOE2 in green, APOE3 in blue and APOE4 in red. Oligo: Oligomycin; FCCP: Carbonyl cyanide-4 (trifluoromethoxy) phenylhydrazone; ROT/AA: Rotenone/Antimycin A. (**E**) Basal respiration, (**F**) maximal respiration, (**G**) spare respiratory capacity (%) and (**H**) H+ leak in iPSCs. (**I**) Basal respiration, **(J**) maximal respiration, (**K**) spare respiratory capacity (%) and (**L**) H+ leak in iN cells. Kruskal-Wallis test (E, F, H) and one-way ANOVA (G, I-L). ns = not significant, * p < 0.05, ** p < 0.01, *** p < 0.001.

## Discussion

In this study, a pure iPSC-derived neuronal culture of APOE-isogenic lines was established to study the role of APOE in neuronal energy metabolism. We demonstrated that APOE2, -E3 and -E4 iN cells express APOE protein of which a small fraction is secreted. This aligns with the literature showing neuronal expression of APOE under physiological conditions [17–20] and only 10 % secretion of neuronally produced APOE [4,6]. In response to injury neuronal APOE expression is upregulated as a potentially protective mechanism, however, upregulation of neuronal APOE4 promoted excitotoxic neuronal cell death [17–20]. APOE4 neurons produce and secrete less total APOE compared to APOE3, but generate more APOE fragments [21]. Neuronal APOE4 is more neurotoxic in the sense of stimulating neuroinflammation by activating microglia and contributing to signaling pathways of the Aβ and tau pathology as well as impairing cholesterol transport by reducing myelin formation [7,22]. However, the exact mechanism by which neuronal APOE4 contributes to AD pathology is still not fully understood. Our study shows that neuronal APOE4 also affects the energy metabolism by increasing mitochondrial and glycolytic ATP production.

As expected, the primary energy source for iN cells in our study was oxidative phosphorylation (OXPHOS), while iPSCs predominantly rely on glycolysis to meet their energy demands. This aligns with earlier studies showing that the process of reprograming into iPSCs results in a glycolytic signature [23]. Neurons predominantly use OXPHOS for ATP production under basal conditions and just switch to glycolysis when facing increased energy needs [24].

Increased ATP production based on glycolysis was observed in APOE4 iN cells and iPSCs. These results support the increase in the glycolytic enzymes that has been observed in the frontal cortex lysate of AD patients as well as glial cells after the treatment with AD plasma [25,26]. Elevated glycolysis has been viewed as a potential compensatory response to mitochondrial dysfunction after exposure to AD plasma, serving as an early indicator of cellular energy deficiency [26,27]. In N2A cells no glycolytic difference between APOE2, -E3 and -E4 cells at early stage could be observed. However, at higher passages APOE4 N2A cells showed impaired glycolysis and glycolytic activity indicating an age-dependent effect of APOE4 on their energy metabolism. This effect could also be observed for hexokinase (HK) expression, which is decreased in APOE4 cells with higher passages whereas APOE2 and APOE3 showed a stable expression of HK over time [28]. Qi and colleagues suggested a cell type-specific APOE effect on glycolysis with showing reduced ATP levels and glycolysis in primary hippocampal neurons and increased glycolytic rates and ATP production in astrocytes [29]. Nonetheless, our study primarily analyzed the glycolytic ATP production, underscoring the need for further research to investigate the impact of APOE4 on neuronal glycolytic metabolism in the future.

We show that APOE regulated mitochondrial respiration and respiratory capacity in a genotype-dependent manner with highest basal and maximal respiration in APOE4 iN cells compared to APOE-KO and highest spare respiratory capacity in APOE2. This aligns with earlier studies showing increased oxidative stress mechanisms and ROS production as well as upregulation of mitochondrial respiratory complexes in iPSC-derived neurons from sporadic AD patients, even in the absence of changes in Aβ or tau and p-tau levels [15]. Mitochondrial and glycolytic impairments had already been linked to AD before [30]. APOE4 carriers show mitochondrial dysfunction in brain areas associated with AD even before the onset of amyloid or tau pathology or cognitive changes indicating that mitochondrial dysfunction is involved in early AD pathology [31]. However, contrary to our findings of mitochondrial upregulation and enhanced ATP production in APOE4 cells, some studies indicated mitochondrial downregulation and decreased ATP production. APOE4 primary hippocampal neurons from mice showed lower maximal respiration and spare respiratory capacity ratio as well as lower mitochondrial membrane potential, reduced expression of complex subunits and reduced ATP levels. [29]. Neuronal APOE4 expression reduced the levels of mitochondrial respiratory complex subunits compared to APOE3 neurons [32,33]. APOE4 expression in N2A cells decreased the enzymatic activity of complex IV [32]. However, these studies have primarily relied on animal models or cell lines, which may have different physiological and metabolic properties compared to human neurons, resulting in different outcomes. Further, Chen and colleagues observed changes in respiratory chain complexes in neurons but not in astrocytes, indicating that mitochondrial dysfunction based on APOE4 expression is neuron-specific [32]. Neuron-specific proteolysis of APOE4, resulting in APOE4 fragments in the cytosol, caused toxic effects such as tau phosphorylation, alterations in cytoskeleton and mitochondrial dysfunctions [34,35].

Mitochondria are highly dynamic organelles undergoing regular fission and fusion cycles. Mitochondrial fusion is particularly important in respiratory active cells, such as neurons [36]. Therefore, we analyzed proteins involved in fusion and fission proteins, but did not observe any APOE-dependent effects in iN cells. This suggests that mitochondrial dynamics are not affected by APOE, independent of further mitochondrial functions including the electron transport chain (ETC). This is in agreement with a previous study showing altered levels of ETC complexes in iN cells from AD patients in the absence of fusion and fission alterations [15], suggesting that human iN cell fusion and fission is a stable mechanism that is not affected by other mitochondrial dysfunctions. In contrast, APOE-dependent alterations in mitochondrial fusion/fission protein levels have been shown in other studies, however those were contradictory. MFN1, MFN2, OPA1 and FIS1 levels were decreased in brains and neurons of AD patients and APOE4 carriers [37,38], whereas APOE4 N2A cells expressed higher levels of fission and fusion proteins than APOE3 N2A cells [33]. These studies suggest that additional mechanisms, beyond the APOE genotype, may play a role in regulating mitochondrial fusion and fission in AD.

In general, many contradictory results on the impact of APOE on energy metabolism have been published. Some studies found that APOE4 reduced glycolysis [28,39], whereas others observed a shift from oxidative to glycolytic metabolism in APOE4 cells with higher glycolytic activity in APOE4 than APOE3 [40–42]. Our study showed enhancement of both, oxidative as well as glycolytic metabolism. The variations may be attributed to discrepancies across studies encompassing differences in cell types, cell species and the time point of analysis.

In addition to APOE4 and APOE3 cells, our study included APOE2 and APOE-KO iN cells. We did not observe differences in intracellular or secreted APOE levels between the isogenic lines but generally higher APOE levels per total protein levels in iPSCs than in iN cells. This indicates that the APOE4-induced increase in neuronal energy metabolism is independent of the levels of APOE4 protein. When comparing APOE-KO cells to the other APOE cell lines, we observed that APOE-KO iN cells have a similar response as APOE2 and APOE3 but not as APOE4, concluding that APOE4 rather displays a gain-of-function than a loss-of-function mechanism. This was also observed in astrocytes from the same iPSC lines, where APOE-KO astrocytes showed a similar response in glutamate and Aβ uptake, cholesterol and lipid metabolism as well as inflammatory response to APOE2 but not to APOE4 astrocytes [14]. This is also consistent with the study of Chemparathy and colleagues showing the protective function of APOE loss-of-variants in healthy individuals and AD patients [43].

Especially the time point of analysis seems to be a critical variable between the studies. On the first view, our data of increased glucose metabolism in APOE4 iN cells contradicts the findings of a reduced metabolism (hypometabolism) observed in APOE4 carrier. However, previous studies have already linked APOE4 to increased (hyper-)metabolism in young individuals. Studies in young adults, before the onset of a pathology, showed that APOE4 carriers display hypermetabolism and hyperactivity in distinct brain regions, such as the hippocampus, entorhinal cortex and cortical regions [44–46]. Further, several studies using APOE4 KI mouse models, that do not yet show features of neurodegeneration, observed a hypermetabolism caused be APOE4 [47,48]. Therefore, we suggest that hypermetabolism is an early feature that may be triggered by APOE4, especially before the onset of AD pathology. However, with age, metabolic activity switches to a hypometabolism. The reason may be that an early hypermetabolism drives AD pathology including amyloid production and aggregation as well as tau phosphorylation and accumulation which leads to neurodegeneration [48]. Subsequently, this neurodegeneration results in a hypoactive state once the disease process has reached a critical stage. As iPSC-models are considered to represent very young cells and are supposed to be a model of early disease or even before disease onset, we observed the early hypermetabolism instead of a hypometabolism. This is further supported by a recent study showing that APP KI mice display mitochondrial hypermetabolism before the onset of any pathology. Upon increasing pathologies, the brain shifted to a state of hypometabolism [49]. In a recent comment, Sercel and colleagues wrote that it was long thought that mitochondrial diseases are linked to ATP deficiency. However, several studies now rather link mitochondrial and OXPHOS deficiency to hypermetabolism [50]. One potential mechanism could be the upregulation of various biological processes and stress responses to compensate for OXPHOS deficits, ultimately resulting in hypermetabolism.

## Conclusions

It is important to recognize that this study only scratches the surface of understanding human neuronal energy metabolism in the context of APOE genotypes. While the observed mitochondrial and glycolytic abnormalities are significant, there is the need for further investigations into the mechanistic basis of the observed APOE4-mediated increase in energy production. Especially the comparison between early and late time points of analysis should be the focus of future research. Further, this study was based on glutamatergic neurons. However, it has been shown that inhibitory neurons express higher levels of APOE4, and the proportion of inhibitory interneurons could be correlated with AD disease progression [4,34,48] highlighting the need for including a wider range of neuronal subtypes in future studies. It will also be interesting to investigate the APOE effect on iPSC-astrocytes from the same isogenic iPSC lines to discriminate between neuronal and astrocytic effects. More studies are necessary to further untangle the molecular pathways and signaling cascades involved in APOE4-induced mitochondrial and glycolytic dysfunction to understand the pathophysiology underlying neurodegenerative diseases such as AD.

## References

1. Scheltens, P.; Strooper, B.D.; Kivipelto, M.; Holstege, H.; Chételat, G.; Teunissen, C.E.; Cummings, J.; van der Flier, W.M. Alzheimer’s Disease. Lancet 2021, 397, 1577–1590, doi:10.1016/S0140-6736(20)32205-4.

2. Andrews, S.J.; Renton, A.E.; Fulton-Howard, B.; Podlesny-Drabiniok, A.; Marcora, E.; Goate, A.M. The Complex Genetic Architecture of Alzheimer’s Disease: Novel Insights and Future Directions. eBioMedicine 2023, 90, 104511, doi:10.1016/j.ebiom.2023.104511.

3. Michaelson, D.M. APOE Epsilon4: The Most Prevalent yet Understudied Risk Factor for Alzheimer’s Disease. Alzheimer’s & dementia : the journal of the Alzheimer’s Association 2014, 10, 861–868, doi:10.1016/j.jalz.2014.06.015.

4. Blumenfeld, J.; Yip, O.; Kim, M.J.; Huang, Y. Cell Type-Specific Roles of APOE4 in Alzheimer Disease. Nat Rev Neurosci 2024, 25, 91–110, doi:10.1038/s41583-023-00776-9.

5. Lanfranco, M.F.; Sepulveda, J.; Kopetsky, G.; Rebeck, G.W. Expression and Secretion of apoE Isoforms in Astrocytes and Microglia during Inflammation. Glia 2021, 69, 1478–1493, doi:10.1002/glia.23974.

6. Harris, F.M.; Tesseur, I.; Brecht, W.J.; Xu, Q.; Mullendorff, K.; Chang, S.; Wyss-Coray, T.; Mahley, R.W.; Huang, Y. Astroglial Regulation of Apolipoprotein E Expression in Neuronal Cells. Implications for Alzheimer’s Disease. J Biol Chem 2004, 279, 3862–3868, doi:10.1074/jbc.M309475200.

7. Zhang, L.; Xia, Y.; Gui, Y. Neuronal ApoE4 in Alzheimer’s Disease and Potential Therapeutic Targets. Front Aging Neurosci 2023, 15, 1199434, doi:10.3389/fnagi.2023.1199434.

8. Yin, F.; Sancheti, H.; Patil, I.; Cadenas, E. Energy Metabolism and Inflammation in Brain Aging and Alzheimer’s Disease. Free radical biology & medicine 2016, 100, 108–122, doi:10.1016/j.freeradbiomed.2016.04.200.

9. An, Y.; Varma, V.R.; Varma, S.; Casanova, R.; Dammer, E.; Pletnikova, O.; Chia, C.W.; Egan, J.M.; Ferrucci, L.; Troncoso, J.; et al. Evidence for Brain Glucose Dysregulation in Alzheimer’s Disease. Alzheimers Dement 2018, 14, 318–329, doi:10.1016/j.jalz.2017.09.011.

10. Ossenkoppele, R.; van der Flier, W.M.; Zwan, M.D.; Adriaanse, S.F.; Boellaard, R.; Windhorst, A.D.; Barkhof, F.; Lammertsma, A.A.; Scheltens, P.; van Berckel, B.N.M. Differential Effect of APOE Genotype on Amyloid Load and Glucose Metabolism in AD Dementia. Neurology 2013, 80, 359–365, doi:10.1212/WNL.0b013e31827f0889.

11. de Leeuw, S.M.; Tackenberg, C. Alzheimer’s in a Dish – Induced Pluripotent Stem Cell-Based Disease Modeling. Translational Neurodegeneration 2019, 8, doi:10.1186/s40035-019-0161-0.

12. Schmid, B.; Prehn, K.R.; Nimsanor, N.; Garcia, B.I.A.; Poulsen, U.; Jørring, I.; Rasmussen, M.A.; Clausen, C.; Mau-Holzmann, U.A.; Ramakrishna, S.; et al. Generation of a Set of Isogenic, Gene-Edited iPSC Lines Homozygous for All Main APOE Variants and an APOE Knock-out Line. Stem Cell Research 2019, 34, 101349, doi:10.1016/j.scr.2018.11.010.

13. Schmid, B.; Prehn, K.R.; Nimsanor, N.; Garcia, B.I.A.; Poulsen, U.; Jørring, I.; Rasmussen, M.A.; Clausen, C.; Mau-Holzmann, U.A.; Ramakrishna, S.; et al. Corrigendum to “Generation of a Set of Isogenic, Gene-Edited iPSC Lines Homozygous for All Main APOE Variants and an APOE Knockout Line” [Stem Cell Res. 34/1873-5061 (2019) 101349-55]. Stem Cell Research 2020, 48, 102005, doi:10.1016/j.scr.2020.102005.

14. de Leeuw, S.M.; Kirschner, A.W.T.; Lindner, K.; Rust, R.; Budny, V.; Wolski, W.E.; Gavin, A.C.; Nitsch, R.M.; Tackenberg, C. APOE2, E3, and E4 Differentially Modulate Cellular Homeostasis, Cholesterol Metabolism, and Inflammatory Response in Isogenic iPSC-Derived Astrocytes. Stem Cell Rep 2022, 17, 110–126, doi:10.1016/j.stemcr.2021.11.007.

15. Birnbaum, J.H.; Wanner, D.; Gietl, A.F.; Saake, A.; Hock, C.; Nitsch, R.M.; Tackenberg, C. Oxidative Stress and Altered Mitochondrial Protein Expression in the Absence of Amyloid-Beta and Tau Pathology in iPSC-Derived Neurons from Sporadic Alzheimer’s Disease Patients. Stem Cell Research 2018, 27, 121–130.

16. Soutschek, M.; Bianco, A.L.; Galkin, S.; Wüst, T.; Colameo, D.; Germade, T.; Gross, F.; Ziegler, L. von; Bohacek, J.; Germain, P.-L.; et al. A Human-Specific microRNA Controls the Timing of Excitatory Synaptogenesis 2023, 2023.10.04.560889.

17. Xu, Q.; Bernardo, A.; Walker, D.; Kanegawa, T.; Mahley, R.W.; Huang, Y. Profile and Regulation of Apolipoprotein E (ApoE) Expression in the CNS in Mice with Targeting of Green Fluorescent Protein Gene to the ApoE Locus. J Neurosci 2006, 26, 4985–4994, doi:10.1523/jneurosci.5476-05.2006.

18. Xu, Q.; Walker, D.; Bernardo, A.; Brodbeck, J.; Balestra, M.E.; Huang, Y. Intron-3 Retention/Splicing Controls Neuronal Expression of Apolipoprotein E in the CNS. J Neurosci 2008, 28, 1452–1459, doi:10.1523/JNEUROSCI.3253-07.2008.

19. Mahley, R.W. Central Nervous System Lipoproteins: ApoE and Regulation of Cholesterol Metabolism. Arteriosclerosis, thrombosis, and vascular biology 2016, 36, 1305–1315, doi:10.1161/atvbaha.116.307023.

20. Buttini, M.; Masliah, E.; Yu, G.-Q.; Palop, J.J.; Chang, S.; Bernardo, A.; Lin, C.; Wyss-Coray, T.; Huang, Y.; Mucke, L. Cellular Source of Apolipoprotein E4 Determines Neuronal Susceptibility to Excitotoxic Injury in Transgenic Mice. Am J Pathol 2010, 177, 563–569, doi:10.2353/ajpath.2010.090973.

21. Wang, C.; Najm, R.; Xu, Q.; Jeong, D.E.; Walker, D.; Balestra, M.E.; Yoon, S.Y.; Yuan, H.; Li, G.; Miller, Z.A.; et al. Gain of Toxic Apolipoprotein E4 Effects in Human iPSC-Derived Neurons Is Ameliorated by a Small-Molecule Structure Corrector. Nat Med 2018, 24, 647–657, doi:10.1038/s41591-018-0004-z.

22. Blanchard, J.W.; Akay, L.A.; Davila-Velderrain, J.; von Maydell, D.; Mathys, H.; Davidson, S.M.; Effenberger, A.; Chen, C.-Y.; Maner-Smith, K.; Hajjar, I.; et al. APOE4 Impairs Myelination via Cholesterol Dysregulation in Oligodendrocytes. Nature 2022, 611, 769–779, doi:10.1038/s41586-022-05439-w.

23. Varum, S.; Rodrigues, A.S.; Moura, M.B.; Momcilovic, O.; Easley, C.A.; Ramalho-Santos, J.; Van Houten, B.; Schatten, G. Energy Metabolism in Human Pluripotent Stem Cells and Their Differentiated Counterparts. PLoS One 2011, 6, e20914, doi:10.1371/journal.pone.0020914.

24. Wei, Y.; Miao, Q.; Zhang, Q.; Mao, S.; Li, M.; Xu, X.; Xia, X.; Wei, K.; Fan, Y.; Zheng, X.; et al. Aerobic Glycolysis Is the Predominant Means of Glucose Metabolism in Neuronal Somata, Which Protects against Oxidative Damage. Nat Neurosci 2023, 26, 2081–2089, doi:10.1038/s41593-023-01476-4.

25. Soucek, T.; Cumming, R.; Dargusch, R.; Maher, P.; Schubert, D. The Regulation of Glucose Metabolism by HIF-1 Mediates a Neuroprotective Response to Amyloid Beta Peptide. Neuron 2003, 39, 43–56, doi:10.1016/s0896-6273(03)00367-2.

26. Jayasena, T.; Poljak, A.; Braidy, N.; Smythe, G.; Raftery, M.; Hill, M.; Brodaty, H.; Trollor, J.; Kochan, N.; Sachdev, P. Upregulation of Glycolytic Enzymes, Mitochondrial Dysfunction and Increased Cytotoxicity in Glial Cells Treated with Alzheimer’s Disease Plasma. PLoS One 2015, 10, e0116092, doi:10.1371/journal.pone.0116092.

27. Zhang, X.; Alshakhshir, N.; Zhao, L. Glycolytic Metabolism, Brain Resilience, and Alzheimer’s Disease. Front Neurosci 2021, 15, 662242, doi:10.3389/fnins.2021.662242.

28. Zhang, X.; Wu, L.; Swerdlow, R.H.; Zhao, L. Opposing Effects of ApoE2 and ApoE4 on Glycolytic Metabolism in Neuronal Aging Supports a Warburg Neuroprotective Cascade against Alzheimer’s Disease. Cells 2023, 12, 410, doi:10.3390/cells12030410.

29. Qi, G.; Mi, Y.; Shi, X.; Gu, H.; Brinton, R.D.; Yin, F. ApoE4 Impairs Neuron-Astrocyte Coupling of Fatty Acid Metabolism. Cell Rep 2021, 34, 108572, doi:10.1016/j.celrep.2020.108572.

30. Terada, T.; Obi, T.; Bunai, T.; Matsudaira, T.; Yoshikawa, E.; Ando, I.; Futatsubashi, M.; Tsukada, H.; Ouchi, Y. In Vivo Mitochondrial and Glycolytic Impairments in Patients with Alzheimer Disease. Neurology 2020, 94, e1592–e1604, doi:10.1212/WNL.0000000000009249.

31. Valla, J.; Yaari, R.; Wolf, A.B.; Kusne, Y.; Beach, T.G.; Roher, A.E.; Corneveaux, J.J.; Huentelman, M.J.; Caselli, R.J.; Reiman, E.M. Reduced Posterior Cingulate Mitochondrial Activity in Expired Young Adult Carriers of the APOE E4 Allele, the Major Late-Onset Alzheimer’s Susceptibility Gene. J Alzheimers Dis 2010, 22, 307–313, doi:10.3233/JAD-2010-100129.

32. Chen, H.-K.; Ji, Z.-S.; Dodson, S.E.; Miranda, R.D.; Rosenblum, C.I.; Reynolds, I.J.; Freedman, S.B.; Weisgraber, K.H.; Huang, Y.; Mahley, R.W. Apolipoprotein E4 Domain Interaction Mediates Detrimental Effects on Mitochondria and Is a Potential Therapeutic Target for Alzheimer Disease. J Biol Chem 2011, 286, 5215–5221, doi:10.1074/jbc.M110.151084.

33. Orr, A.L.; Kim, C.; Jimenez-Morales, D.; Newton, B.W.; Johnson, J.R.; Krogan, N.J.; Swaney, D.L.; Mahley, R.W. Neuronal Apolipoprotein E4 Expression Results in Proteome-Wide Alterations and Compromises Bioenergetic Capacity by Disrupting Mitochondrial Function. J Alzheimers Dis 2019, 68, 991–1011, doi:10.3233/JAD-181184.

34. Mahley, R.W. Apolipoprotein E4 Targets Mitochondria and the Mitochondria-Associated Membrane Complex in Neuropathology, Including Alzheimer’s Disease. Curr Opin Neurobiol 2023, 79, 102684, doi:10.1016/j.conb.2023.102684.

35. Mahley, R.W.; Huang, Y. Apolipoprotein e Sets the Stage: Response to Injury Triggers Neuropathology. Neuron 2012, 76, 871–885, doi:10.1016/j.neuron.2012.11.020.

36. Westermann, B. Bioenergetic Role of Mitochondrial Fusion and Fission. Biochimica et Biophysica Acta (BBA) - Bioenergetics 2012, 1817, 1833–1838, doi:10.1016/j.bbabio.2012.02.033.

37. Wang, X.; Su, B.; Fujioka, H.; Zhu, X. Dynamin-like Protein 1 Reduction Underlies Mitochondrial Morphology and Distribution Abnormalities in Fibroblasts from Sporadic Alzheimer’s Disease Patients. Am J Pathol 2008, 173, 470–482, doi:10.2353/ajpath.2008.071208.

38. Yin, J.; Reiman, E.M.; Beach, T.G.; Serrano, G.E.; Sabbagh, M.N.; Nielsen, M.; Caselli, R.J.; Shi, J. Effect of ApoE Isoforms on Mitochondria in Alzheimer Disease. Neurology 2020, 94, e2404–e2411, doi:10.1212/WNL.0000000000009582.

39. Fang, W.; Xiao, N.; Zeng, G.; Bi, D.; Dai, X.; Mi, X.; Ye, Q.; Chen, X.; Zhang, J. APOE4 Genotype Exacerbates the Depression-like Behavior of Mice during Aging through ATP Decline. Transl Psychiatry 2021, 11, 507, doi:10.1038/s41398-021-01631-0.

40. Lee, H.; Cho, S.; Kim, M.-J.; Park, Y.J.; Cho, E.; Jo, Y.S.; Kim, Y.-S.; Lee, J.Y.; Thoudam, T.; Woo, S.-H.; et al. ApoE4-Dependent Lysosomal Cholesterol Accumulation Impairs Mitochondrial Homeostasis and Oxidative Phosphorylation in Human Astrocytes. Cell Rep 2023, 42, 113183, doi:10.1016/j.celrep.2023.113183.

41. Farmer, B.C.; Williams, H.C.; Devanney, N.A.; Piron, M.A.; Nation, G.K.; Carter, D.J.; Walsh, A.E.; Khanal, R.; Young, L.E.A.; Kluemper, J.C.; et al. APOE4 Lowers Energy Expenditure in Females and Impairs Glucose Oxidation by Increasing Flux through Aerobic Glycolysis. Molecular neurodegeneration 2021, 16, 62, doi:10.1186/s13024-021-00483-y.

42. Sonntag, K.C.; Ryu, W.I.; Amirault, K.M.; Healy, R.A.; Siegel, A.J.; McPhie, D.L.; Forester, B.; Cohen, B.M. Late-Onset Alzheimer’s Disease Is Associated with Inherent Changes in Bioenergetics Profiles. Scientific reports 2017, 7, 14038, doi:10.1038/s41598-017-14420-x.

43. Chemparathy, A.; Le Guen, Y.; Chen, S.; Lee, E.-G.; Leong, L.; Gorzynski, J.E.; Jensen, T.D.; Ferrasse, A.; Xu, G.; Xiang, H.; et al. APOE Loss-of-Function Variants: Compatible with Longevity and Associated with Resistance to Alzheimer’s Disease Pathology. Neuron 2024, 112, 1110-1116.e5, doi:10.1016/j.neuron.2024.01.008.

44. Dennis, N.A.; Browndyke, J.N.; Stokes, J.; Need, A.; Burke, J.R.; Welsh-Bohmer, K.A.; Cabeza, R. Temporal Lobe Functional Activity and Connectivity in Young Adult APOE Varepsilon4 Carriers. Alzheimers Dement 2010, 6, 303–311, doi:10.1016/j.jalz.2009.07.003.

45. Filippini, N.; MacIntosh, B.J.; Hough, M.G.; Goodwin, G.M.; Frisoni, G.B.; Smith, S.M.; Matthews, P.M.; Beckmann, C.F.; Mackay, C.E. Distinct Patterns of Brain Activity in Young Carriers of the APOE-Epsilon4 Allele. Proc Natl Acad Sci U S A 2009, 106, 7209–7214, doi:10.1073/pnas.0811879106.

46. Bookheimer, S.Y.; Strojwas, M.H.; Cohen, M.S.; Saunders, A.M.; Pericak-Vance, M.A.; Mazziotta, J.C.; Small, G.W. Patterns of Brain Activation in People at Risk for Alzheimer’s Disease. N Engl J Med 2000, 343, 450–456, doi:10.1056/NEJM200008173430701.

47. Venzi, M.; Tóth, M.; Häggkvist, J.; Bogstedt, A.; Rachalski, A.; Mattsson, A.; Frumento, P.; Farde, Differential Effect of APOE Alleles on Brain Glucose Metabolism in Targeted Replacement Mice: An [18F]FDG-μPET Study. J Alzheimers Dis Rep 2017, 1, 169–180, doi:10.3233/ADR-170006.

48. Nuriel, T.; Angulo, S.L.; Khan, U.; Ashok, A.; Chen, Q.; Figueroa, H.Y.; Emrani, S.; Liu, L.; Herman, M.; Barrett, G.; et al. Neuronal Hyperactivity Due to Loss of Inhibitory Tone in APOE4 Mice Lacking Alzheimer’s Disease-like Pathology. Nat Commun 2017, 8, 1464, doi:10.1038/s41467-017-01444-0.

49. Naia, L.; Shimozawa, M.; Bereczki, E.; Li, X.; Liu, J.; Jiang, R.; Giraud, R.; Leal, N.S.; Pinho, C.M.; Berger, E.; et al. Mitochondrial Hypermetabolism Precedes Impaired Autophagy and Synaptic Disorganization in App Knock-in Alzheimer Mouse Models. Mol Psychiatry 2023, 28, 3966–3981, doi:10.1038/s41380-023-02289-4.

50. Sercel, A.J.; Sturm, G.; Gallagher, D.; St-Onge, M.-P.; Kempes, C.P.; Pontzer, H.; Hirano, M.; Picard, M. Hypermetabolism and Energetic Constraints in Mitochondrial Disorders. Nat Metab 2024, 6, 192–195, doi:10.1038/s42255-023-00968-8.

