## Supplementary figures for "APOE4 increases energy metabolism in APOE-isogenic iPSC-derived neurons"

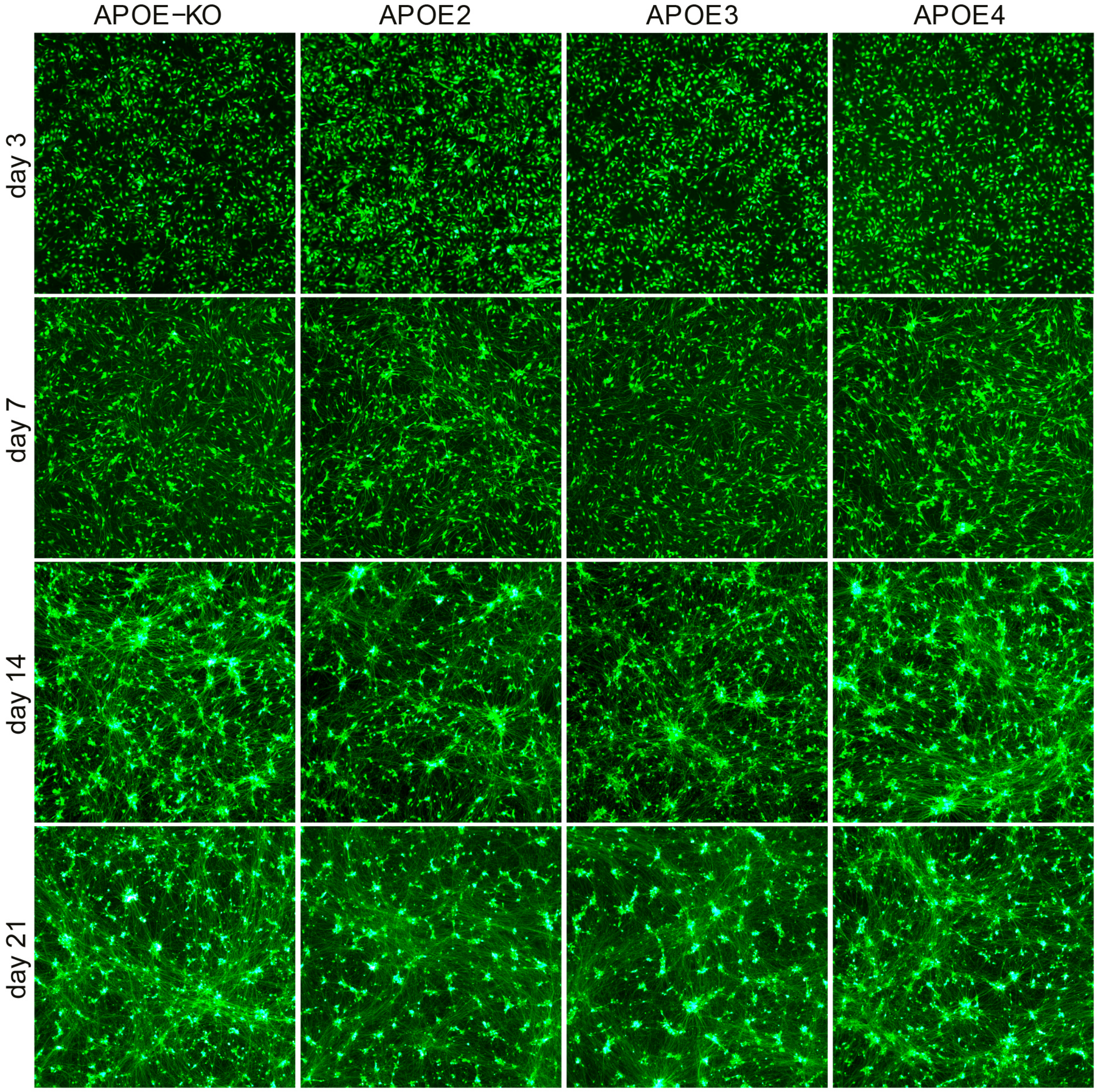


**Figure S1: iN cell differentiation over time.** Representative fluorescence images of GFP-expressing APOE-isogenic iN cells at different time points after induction of differentiation. Scale bar: 50μm

**Table S1: List of qPCR primer.**

| | | |
|---|---|---|
| **GAPDH** | FWD | ACC ACA GTC CAT GCC ATC AC |
| | REV | TCC ACC CTG TTG CTG TA |
| **MAP2** | FWD | CTC AGC ACC GCT AAC AGA GG |
| | REV | CAT TGG CGC TTC GGA CAA G |
| **OCT4** | FWD | GAG AAG CTG GAG CAA AAC CC |
| | REV | ACC TTC CCA AAT AGA ACC CCC |
| **TUBB3** | FWD | GGC CAA GGG TCA CTA CAC G |
| | REV | GCA GTC GCA GTT TTC ACA CTC |
